# Liver sinusoidal endothelial cells orchestrate NK cell recruitment and activation in acute inflammatory liver injury

**DOI:** 10.1101/2022.07.15.500206

**Authors:** Sophia Papaioannou, Jia-Xiang See, Mingeum Jeong, Carolina De La Torre, Philipp-Sebastian Reiners-Koch, Ankita Sati, Carolin Mogler, Michael Platten, Adelheid Cerwenka, Ana Stojanovic

**Author notes:** Equally contributing first authors (Sophia Papaioannou, Jia-Xiang See). Equally contributing senior authors (Adelheid Cerwenka, Ana Stojanovic). **Corresponding Authors:** Ana Stojanovic and Adelheid Cerwenka.

## Abstract

In both steady-state and during endotoxicosis, liver sinusoidal endothelial cells (LSECs) can rapidly clear lipopolysaccharide (LPS) from the bloodstream. They are located along blood sinusoids of the liver, and establish intimate contact with circulating and tissue-resident immune cells. However, their role in regulating immune responses during LPS-induced endotoxicosis remains poorly understood. Here, we show that LSECs play a dual role in regulating inflammatory responses, acting as modulators of NK cell pro-inflammatory output and as major producers of immune cell-attracting chemokines. We demonstrate that LSECs switch their chemokine expression pattern driven by LPS and IFN-γ, resulting in the production of the myeloid-attracting chemokine CCL2 and the lymphoid-attracting chemokine CXCL10, which accumulate in the serum of LPS-challenged animals. In livers of LPS-injected mice, monocytes and Kupffer cells expressed highest amounts of the pro-inflammatory cytokine *Il12a* and *Il18* transcripts, while NK cells expressed the highest amounts of *Ifng*. NK cell exposure to LSECs *in vitro* led to global transcriptomic changes, and primed NK cells to produce higher amounts of IFN-γ in response to IL-12 and IL-18. LSECs required exposure to IFN-γ for *Cxcl10* expression, and *Cxcl10* gene-deletion in endothelial cells abrogated NK cell accumulation in the liver after LPS treatment. Thus, our data indicate that LSECs occupy a unique temporal and spatial position acting as central regulators that respond to both LPS and immune-derived inflammatory signals, and fuel a positive feedback loop of immune cell attraction and activation in the inflamed liver tissue.

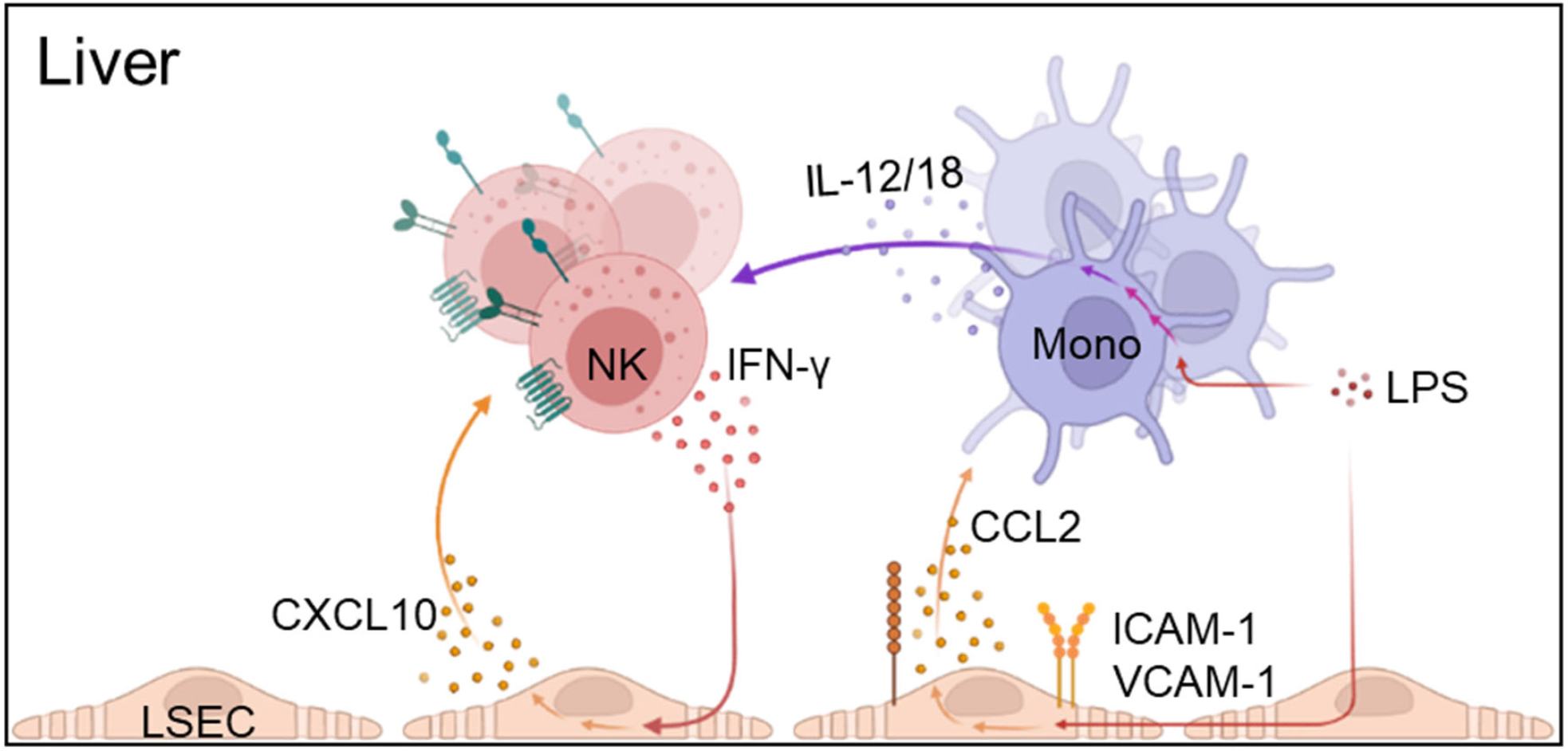

## Introduction

The liver vasculature comprises ramified blood vessels, the sinusoids, lined by specialized liver sinusoidal endothelial cells (LSECs)(Knolle and Wohlleber, 2016). They form highly permeable capillary vessels without a basement membrane. Due to the low blood flow rate and the expression of scavenging receptors, LSECs efficiently endocytose blood-borne pathogens and molecules entering the liver via the hepatic artery and the portal vein (Ganesan et al., 2012; Ganesan et al., 2011). In homeostatic conditions, LSECs were shown to rapidly eliminate the gram-negative bacterial product lipopolysaccharide (LPS) from the circulation, without causing inflammation (Uhrig et al., 2005). However, during bacterial infections, elevated concentrations of LPS in the blood can activate LSECs via Toll-like receptor 4 (TLR4) (Yao et al., 2016), leading to the secretion of pro-inflammatory cytokines and increased expression of adhesion molecules (Cabral et al., 2021; Uhrig et al., 2005). The endothelial response is an integral component of inflammation that regulates immune cell trafficking and functions (Amersfoort et al., 2022; Ducimetiere et al., 2021), but the exact mechanisms and relative contribution of endothelium to inflammatory output in different organs and in different pathological settings are not yet fully understood.

The liver is enriched in innate lymphocytes, mainly comprising circulating Natural Killer (NK) cells and liver-resident type 1 innate lymphoid cells (ILC1s). Both liver-resident and circulatory lymphocytes can continuously interact with LSECs within liver sinusoids. NK cells and ILC1s provide the early defense against infections and malignancies by exerting cytotoxicity and by secreting pro-inflammatory cytokines (Ducimetiere et al., 2021; Weizman et al., 2017). NK cells were shown to sustain LPS-induced inflammation via TNF-α and IFN-γ secretion (Chan et al., 2014), and their depletion led to improved survival of animals upon treatment with LPS. Similarly, in a shock-syndrome model induced by LPS and 2-amino-2-deoxy-D-galactose (D-gal) injection, the depletion of NK cells reduced the mortality of treated animals (Vinay et al., 2004). Thus, NK cells act as crucial effector cells in acute liver disease mediating tissue damage and inflammation. However, the local tissue microenvironment that supports NK cell functions in inflamed liver remains not fully explored.

Here, we demonstrate that LSECs are the major producers of chemokines that accumulate in the serum and attract immune cells in the liver tissue upon LPS-induced acute inflammation. The accumulation of activated NK cells in inflamed liver tissue is supported by endothelium-derived CXCL10. CXCL10 expression by LSECs required exposure to IFN-γ, which is mainly produced by the NK cells and ILCs in the liver upon LPS treatment. We show that the interaction with LSECs leads to NK cell transcriptional reprogramming, and results in priming of NK cells for the enhanced production of IFN-γ, thus supporting the amplification of the inflammatory response in the liver.

## Results

### LSECs are the main producers of chemokines upon LPS treatment

Systemic exposure to LPS induces an accumulation of activated immune cells in the liver tissue driving acute liver inflammation (Sun et al., 2022). Accordingly, mice injected with LPS displayed reduced body and liver weight, and a reduced liver-to-body weight ratio compared to control animals (Figure 1A). The absolute numbers of neutrophils, monocytes, Kupffer cells, as well as NK cells, increased in the livers of LPS-treated mice (Figure 1B-C), while numbers of ILC1s and T cells remained unchanged (Figure 1D and Supplemental Figure 1A). In inflamed livers, increased expression of CD25, CD69, CD11c and Ly6C by NK cells indicated an activated phenotype (Figure 1E-F). However, the expression of Ki67 remained unchanged (Supplemental Figure 1B), suggesting that NK cells did not undergo a change in their proliferative behavior and that active recruitment might contribute to the increased NK cell numbers in the liver tissue after LPS injection. Accordingly, we detected reduced NK cell numbers in the spleen of LPS-injected mice (Supplemental Figure 1C), implicating that NK cells might migrate from the spleen to the liver.

**Figure 1.**
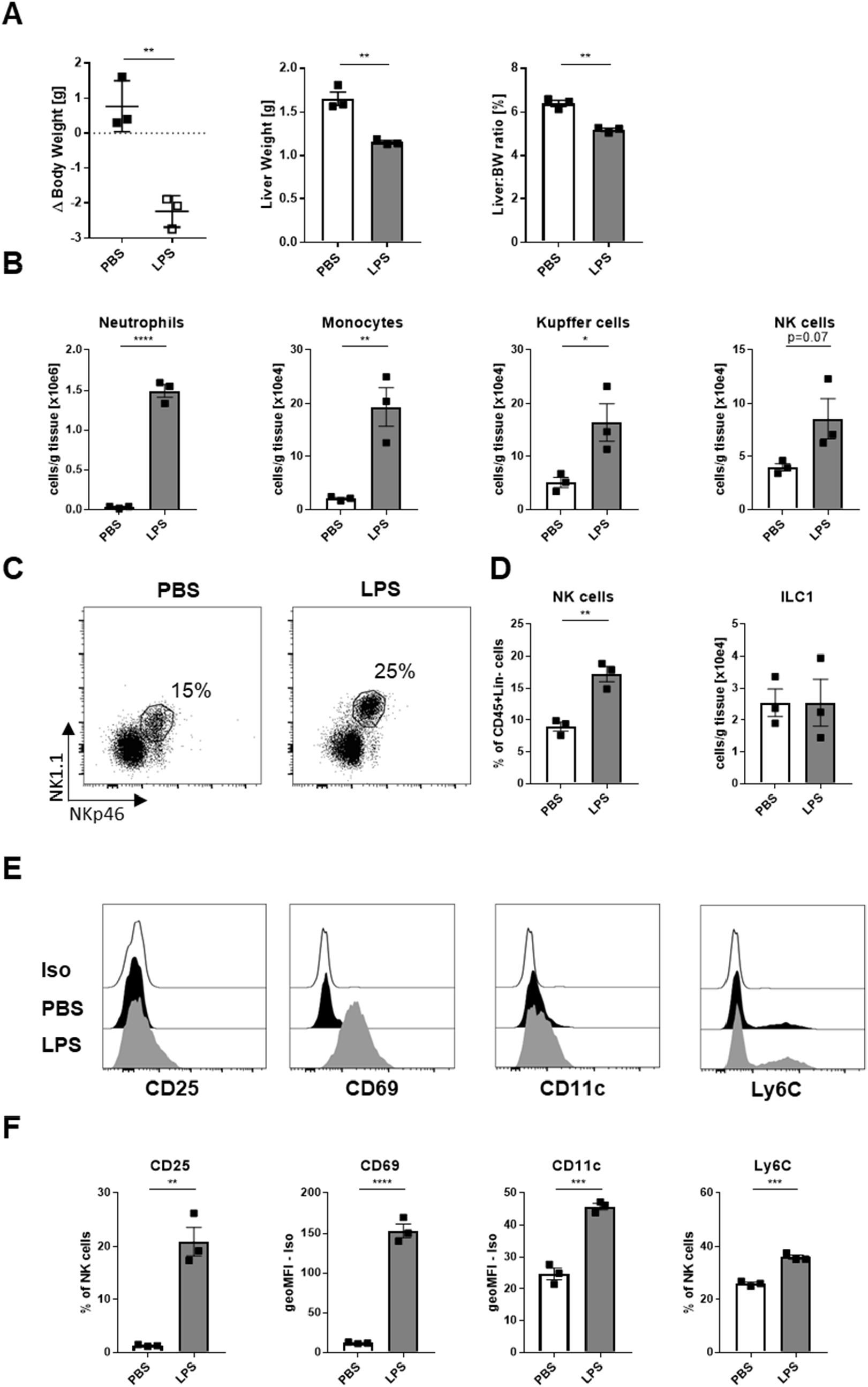
Activated NK cells accumulate in the livers of LPS-treated mice. (A) Body weight after treatment relative to initial body weight (left), liver weight (middle) and liver to body weight (BW) ratio. (B) Cell numbers of indicated immune cell subsets 16 h post-injection of PBS or LPS. (C) Representative flow cytometry dot plots of gated live CD45^+^ Neutrophils (Ly6G^+^), Monocytes (Ly6G^neg^CD11b^+^F4/80^dim^), Kupffer cells (Ly6G^neg^CD11b^dim^F4/80^+^) and NK cells (Ly6G^neg^CD19^neg^ CD3ε^neg^ NK1.1^+^NKp46^+^Eomes^+^) in the liver of PBS- and LPS-injected mice. (D) Frequency of NK cells (NK1.1^+^NKp46^+^Eomes^+^) and ILC1s (NK1.1^+^NKp46^+^Eomes^neg^) among CD45^+^ Lineage (Ly6G, CD19, CD3ε)^neg^ cells in the liver 16 h post-injection of PBS or LPS. (E+F) Representative flow-cytometry histograms (E) and (F) graphs showing expression or frequencies of CD25-, CD69-, CD11c- and Ly6C-expressing NK cells in the livers of PBS- or LPS-injected mice 16 h post-injection. (A-D, F) Data are shown as mean ± SEM, analyzed by unpaired Student’s t-Test. *, p<0.05, **, p<0.01, ****, p<0.0001. Dots represent individual mice. Iso, Isotype; geoMFI, geometric mean fluorescent intensity.

To determine if chemokine gradients support the NK cell recruitment to inflamed liver tissue, we first analyzed the serum of affected animals. We detected elevated concentrations of CCL2, CCL3, CCL4, CCL5, CCL7 and CXCL10 in LPS-treated mice compared to controls (Figure 2A). In particular, CXCL10 concentration was elevated about 55 times in the serum of mice treated with LPS compared to controls. CXCL10 binds to the chemokine receptor CXCR3 (Loetscher et al., 1996). Accordingly, a subset of NK cells expressed CXCR3 in the liver and the spleen of both control and LPS-injected mice (Figure 2B and 2C). To dissect the cellular sources of the chemokines accumulating in the serum, we assessed chemokine production by both immune and non-parenchymal cells from livers, including LSECs, monocytes and Kupffer cells, T cells, and NK cells and ILC1s (Supplemental Figure 2). In inflamed livers, LSECs expressed transcripts encoding all reported CXCR3-ligands, *Cxcl9, Cxcl10* and *Cxcl11* (Figure 3A), and the highest relative amount of *Cxcl10* mRNA among analyzed cell populations (Figure 3B left). Furthermore, LSECs also expressed transcript encoding CCL2 (Figure 3A and 3B right), which was shown to be crucial for the recruitment of inflammatory CCR2-expressing monocytes to the inflamed or damaged livers (Mossanen et al., 2016). These observations correlated with reduced numbers of endothelial cells with “classical” LSEC phenotype, defined by expression of CD146, CD31 and Stabilin2, in LPS-injected compared to control animals (Figure 3C), as well as with higher expression of ICAM-1 (Figure 3D) and VCAM-1 (Figure 3E) on LSECs in inflamed livers, indicative of endothelial cell activation in inflammatory settings (Geraud et al., 2017; Geraud et al., 2013; Harjunpaa et al., 2019). Thus, upon LPS challenge, activated LSECs are the main producers of chemokines in the liver with the potential to shape the liver immune environment.

**Figure 2.**
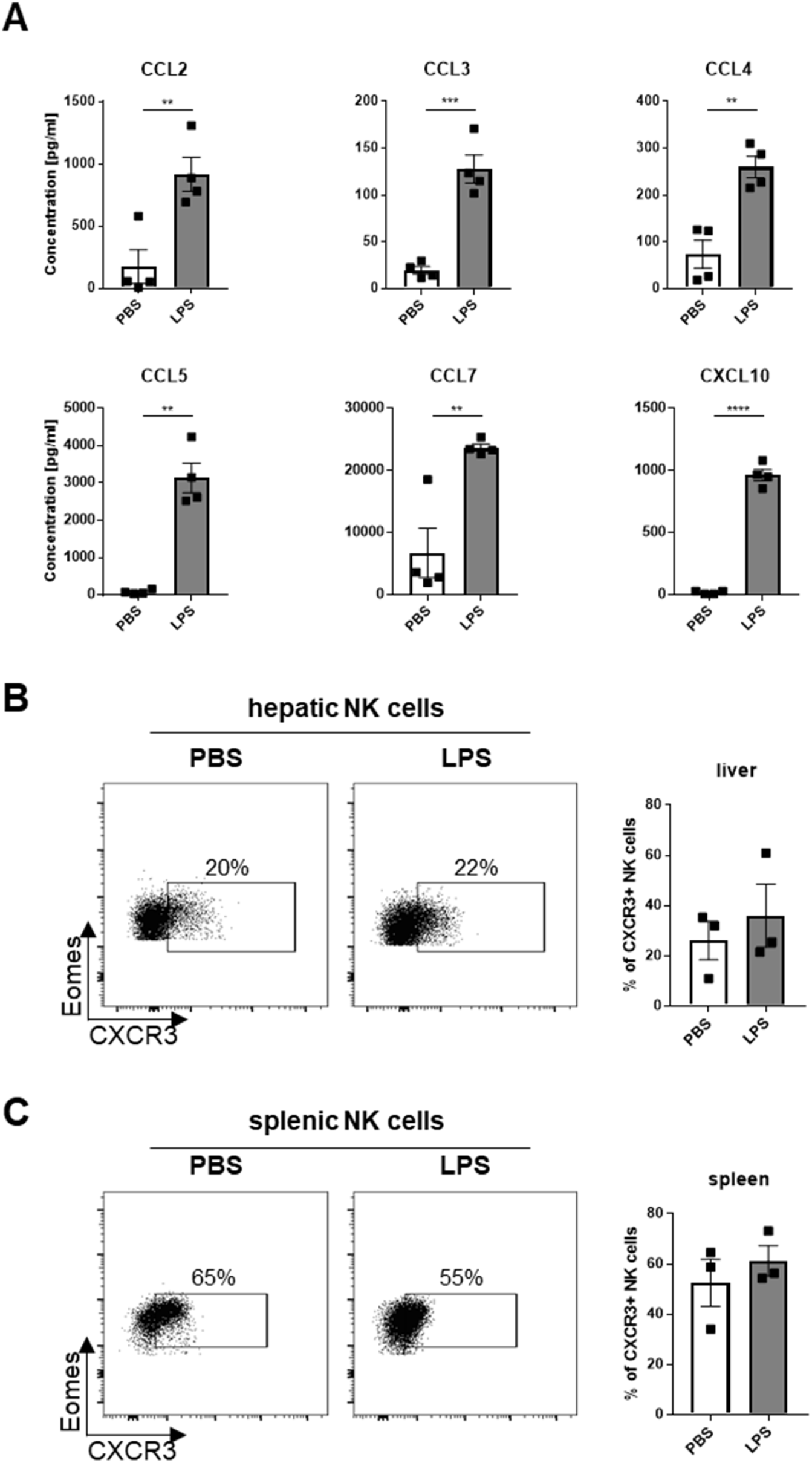
Increased CXCL10 concentration in serum and CXCR3 expression by hepatic and splenic NK cells in LPS-treated mice. (A) Concentration of depicted chemokines in the serum of PBS- and LPS-injected mice 16 h post-injection, assessed by bead-based flow cytometry analysis. (B-C) Representative flow cytometry dot plots and graphs showing the frequency of CXCR3^+^ NK cells (NK1.1^+^NKp46^+^Eomes^+^) in the liver (B) and spleen (C). Data are shown as mean ± SEM, analyzed by unpaired Student’s t-Test. **, p<0.01, ***, p<0.001, ****, p<0.0001. Dots represent individual mice.

**Figure 3.**
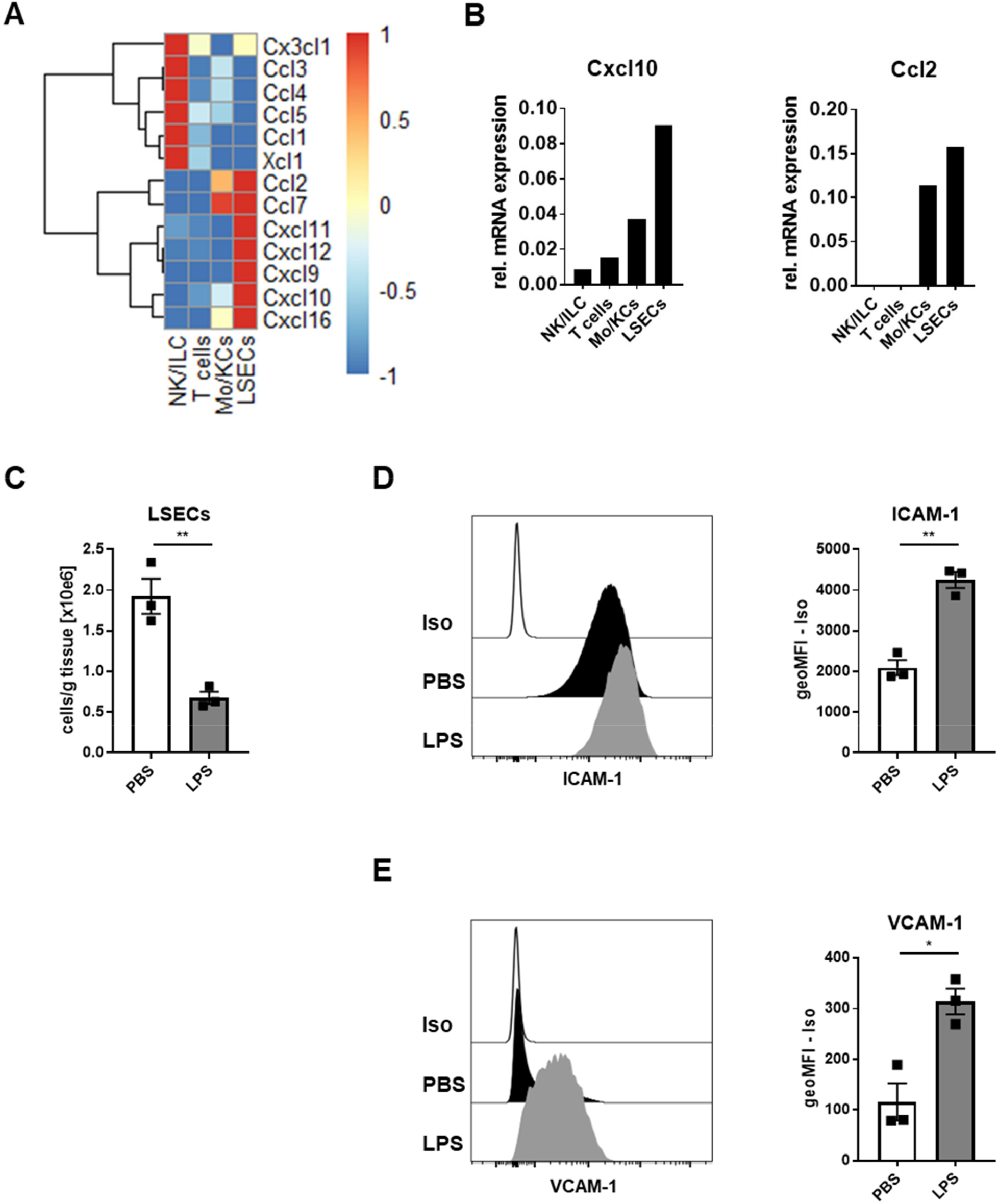
Activated LSECs are the major source of CXCL10 in the liver. (A) Mice were challenged with LPS, and 16 h later LSECs, NK1.1^+^NKp46^+^ cells (NK/ILC), T cells and CD11b^+^F4/80^+^ cells (Mo/KCs) were sorted from 4 pooled liver tissue samples. Heatmap displaying the relative mRNA expression of depicted chemokines in the sorted cell populations, analyzed by qPCR. Data are scaled per row. (B) Relative mRNA expression of *Cxcl10* and *Ccl2* by NK/ILC, T cells, Mo/KCs and LSECs. (C) Cell numbers and (D-E) representative histograms and graphs showing the geometric mean fluorescence intensity (geoMFI) of ICAM-1 (D) and VCAM-1 (E) expression by LSECs (gated as live CD45^neg^Ly6G^neg^CD146^+^CD31^+^Stab2^+^ cells) in PBS- and LPS-injected mice. (C-E) Data are shown as mean ± SEM, analyzed by unpaired Student’s t-Test. *, p<0.05, **, p<0.01. Dots represent individual mice. Iso, Isotype.

### LPS and IFN-γ differentially regulate chemokine expression of LSECs

To determine whether LPS directly affected chemokine expression by LSECs, we isolated LSECs from healthy animals and exposed them to LPS *in vitro*. The treatment induced the transcription of *Ccl2, Ccl3, Ccl4, Ccl5* and *Ccl7*, but not *Cxcl10* (Figure 4A-C). Previous report demonstrated that *Cxcl10* expression can be induced by IFN-γ-receptor engagement (Luster and Ravetch, 1987). Accordingly, exposure of LSECs to IFN-*γ in vitro* increased the expression of *Cxcl10*, as well as the transcripts of the other CXCR3-ligands, *Cxcl9* and *Cxcl11* (Figure 4C). Unlike *Cxcl10*, the expression of VCAM-1 and ICAM-1 can be elevated on LSECs by both LPS and IFN-γ (Figure 4D). These data indicate that LPS and IFN-γ have complementary, non-overlapping roles in inducing the LSEC inflammatory phenotype, by shaping the chemokine profile of the inflamed liver endothelium.

**Figure 4.**
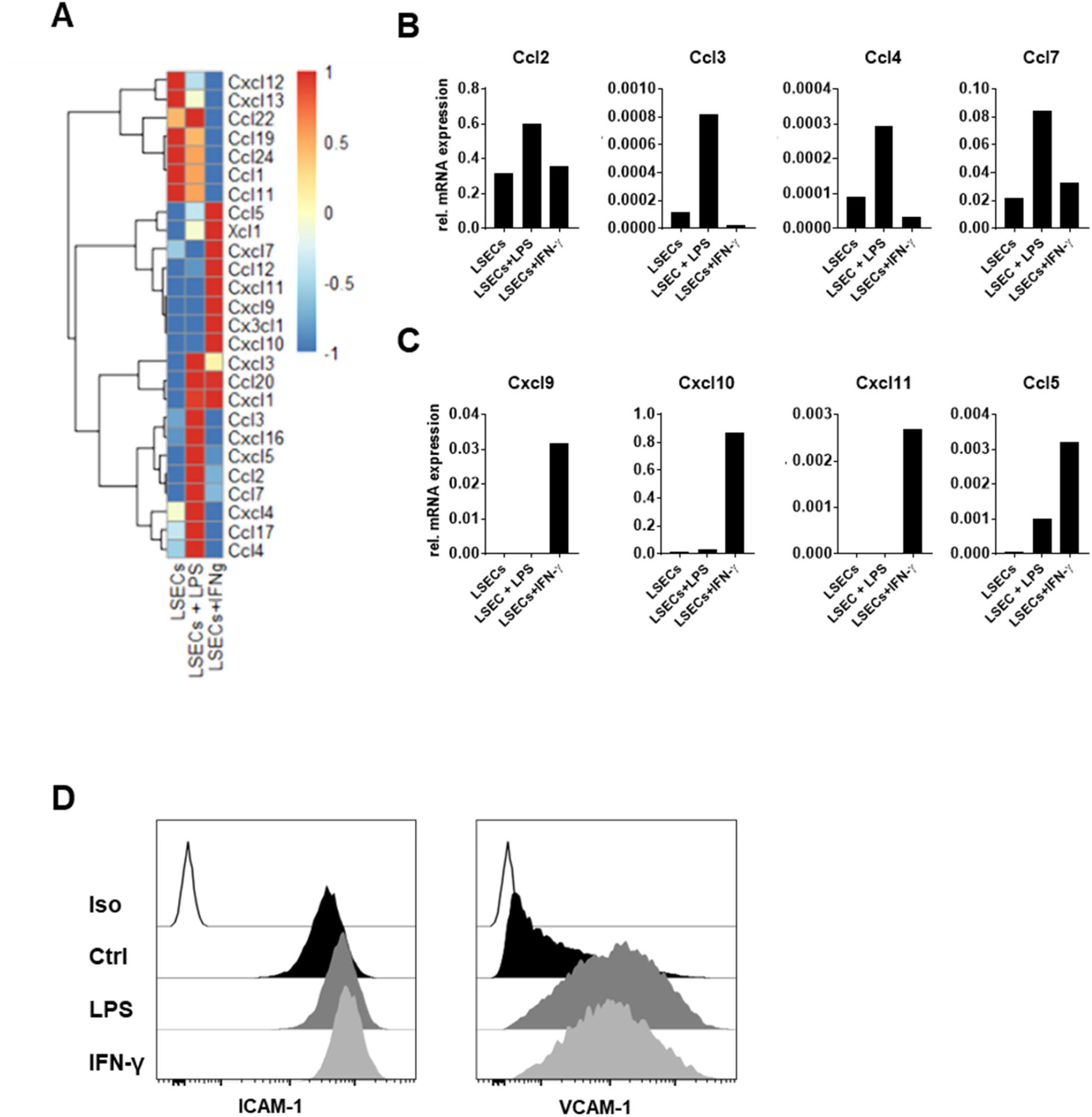
LPS and IFN-γ shape the chemokine profile of LSECs. LSECs purified from murine livers were treated with LPS or IFN-γ for 24h. (A) Heatmap displaying the relative mRNA expression of depicted chemokines, analyzed by qPCR. Data are scaled per row. (B-C) Relative mRNA expression of selected chemokines. (D) Representative histograms of ICAM-1 and VCAM-1 expression on LSECs cultured alone (Ctrl) or stimulated with LPS or IFN-γ. Iso, Isotype.

### Liver NK cells and ILC1s are the major producers of IFN-γ upon LPS challenge

Given the role of IFN-γ in shaping chemokine production by LSECs, we aimed to identify the IFN-γ-producing cells in the liver of LPS-challenged mice. We observed that in LPS-treated mice, IFN-γ was mostly produced by NK1.1^+^NKp46^+^ innate lymphocytes (comprising NK cells and ILC1s), and to a lower extent by T cells (Figure 5A-B). IFN-γ production by NK cells can be triggered by activating receptors via the engagement of ligands, expressed on target cells, or by soluble inflammatory mediators that engage cytokine receptors expressed by NK cells (Long et al., 2013; Mujal et al., 2021). IL-12 and IL-18 are shown to synergistically induce IFN-γ production in NK cells (Fehniger et al., 1999; Lusty et al., 2017). In the liver, monocytes and Kupffer cells were the main producers of these cytokines (Figure 5A-B). To investigate if LSECs affected IFN-γ production by NK cells, we co-cultured LSECs with freshly isolated liver NK cells *in vitro*. While LSECs themselves did not induce IFN-γ production by NK cells (Supplemental Figure 3), they significantly upregulated the NK cell response to IL-12 and IL-18 (Figure 5C). These data indicate that exposure to LSECs shapes NK cell reactivity in inflamed tissue. Accordingly, transcriptional analysis of NK cells co-cultured with LSECs led to significant changes in NK cell transcriptome, affecting the pathways related to cytokine production, leukocyte migration and cell-to-cell adhesion (Figure 5D-E). These include transcripts encoding L-selectin, CD62L (*Sell*) and the chemokine receptor CX3CR1, as well as transcription factors *Zbtb32* and *Irf8*, shown to support NK cell expansion in response to viral infection (Adams et al., 2018; Beaulieu et al., 2014), and *Slc7a5*, encoding system L-amino acid transporter expressed predominantly by activated NK cells (Loftus et al., 2018). In contrast, the expression of *Cd96*, reported to limit NK cell IFN-γ production in response to LPS (Chan et al., 2014), was reduced. Thus, in LPS-induced liver inflammation, LSEC-driven signals affect NK cell responsiveness to pro-inflammatory cytokines provided by accessory myeloid cells, and in turn, NK cell-derived IFN-γ facilitates LSEC chemokine production.

**Figure 5.**
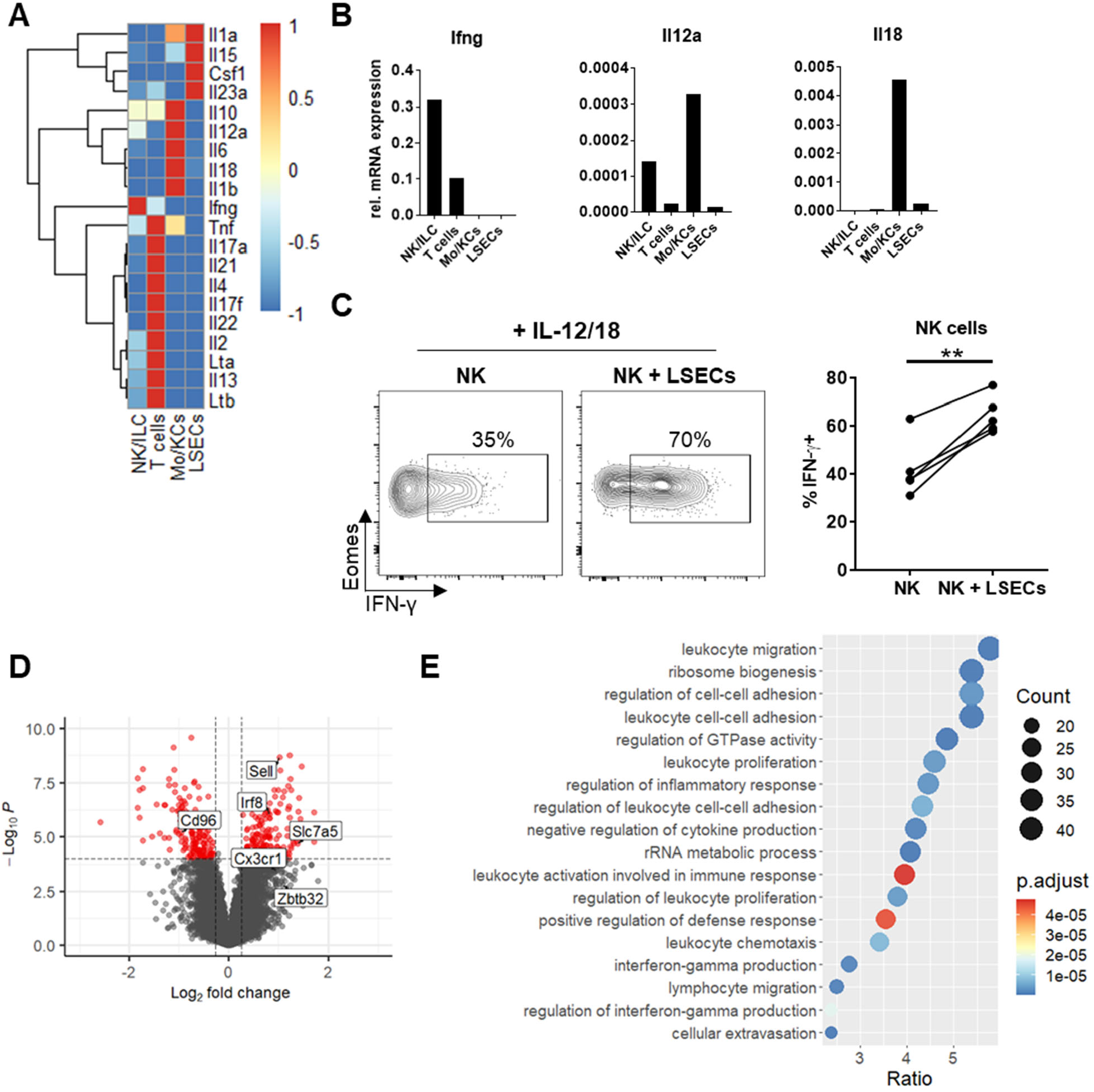
LSECs induce transcriptional changes in NK cells and enhance their IFN-γ production in the presence of IL-12/18. (A) Mice were challenged with LPS, and 16 h later LSECs, NK1.1^+^NKp46^+^ cells (NK/ILC), T cells and CD11b^+^F4/80^+^ cells (Mo/KCs) were sorted from 4 pooled liver tissue samples. Heatmap displaying the relative mRNA expression of depicted cytokines in the sorted cell populations, analyzed by qPCR. Data are scaled per row. (B) Relative mRNA expression of *Ifng, Il12a* and *Il18* in the sorted cell populations. (C-E) Sorted NK cells and purified LSECs were co-cultured overnight, followed by 5 h stimulation with IL-12 and IL-18. (C) Representative flow cytometry contour plots and graph showing the frequency of IFN-γ-expressing NK cells after co-culture and stimulation. Data are shown as mean ± SEM, analyzed by unpaired Student’s t-Test. Each dot represents an independent experiment (n = 4). **, p<0.01. (D) Volcano-plot showing differentially expressed transcripts in NK cells after co-culture with LSECs in red (cut-off p-value (P) = 0.0001, Fold Change (FC) = 1.2). (E) Gene ontology (GO)-enrichment analysis of upregulated transcripts in NK cells co-cultured with LSECs. Selected pathways are depicted. ES, enrichment score; NES, normalized enrichment score, p.adjust, adjusted p-value.

### LSEC-derived CXCL10 is crucial for NK cell recruitment to inflamed liver tissue

To address the contribution of LSEC-derived CXCL10 to NK cell accumulation in inflamed liver tissue *in vivo*, we analyzed mice with an endothelial-specific deficiency of *Cxcl10* (Cdh5^Cre^Cxcl10^fl/fl^). Untreated Cdh5^Cre^Cxcl10^fl/fl^ mice showed no differences in NK cell frequency or cell numbers compared to Cxcl10^fl/fl^ littermates (Figure 6A). Upon challenge with LPS, LSECs expressed similar amounts of VCAM-1 and ICAM-1 irrespective of genotype, indicating comparable endothelial activation in Cdh5^Cre^Cxcl10^fl/fl^ and control animals (Figure 6B). Furthermore, endothelial *Cxcl10* deficiency had no effect on the recruitment of neutrophils and inflammatory monocytes (Figure 6C). However, while NK cells accumulated in the livers of Cxcl10^fl/fl^ mice upon LPS injection, LPS treatment did not lead to a significant increase in NK cell numbers in Cdh5^Cre^Cxcl10^fl/fl^ mice, when compared to PBS-injected controls (Figure 6D). We observed no difference regarding CD25 and CD69 upregulation on liver NK cells from Cxcl10^fl/fl^ and Cdh5^Cre^Cxcl10^fl/fl^ mice after LPS treatment (Figure 6E). These data indicate that endothelium-derived CXCL10 is a major driver of NK cell migration to the inflamed liver tissue, without affecting NK cell activation.

**Figure 6.**
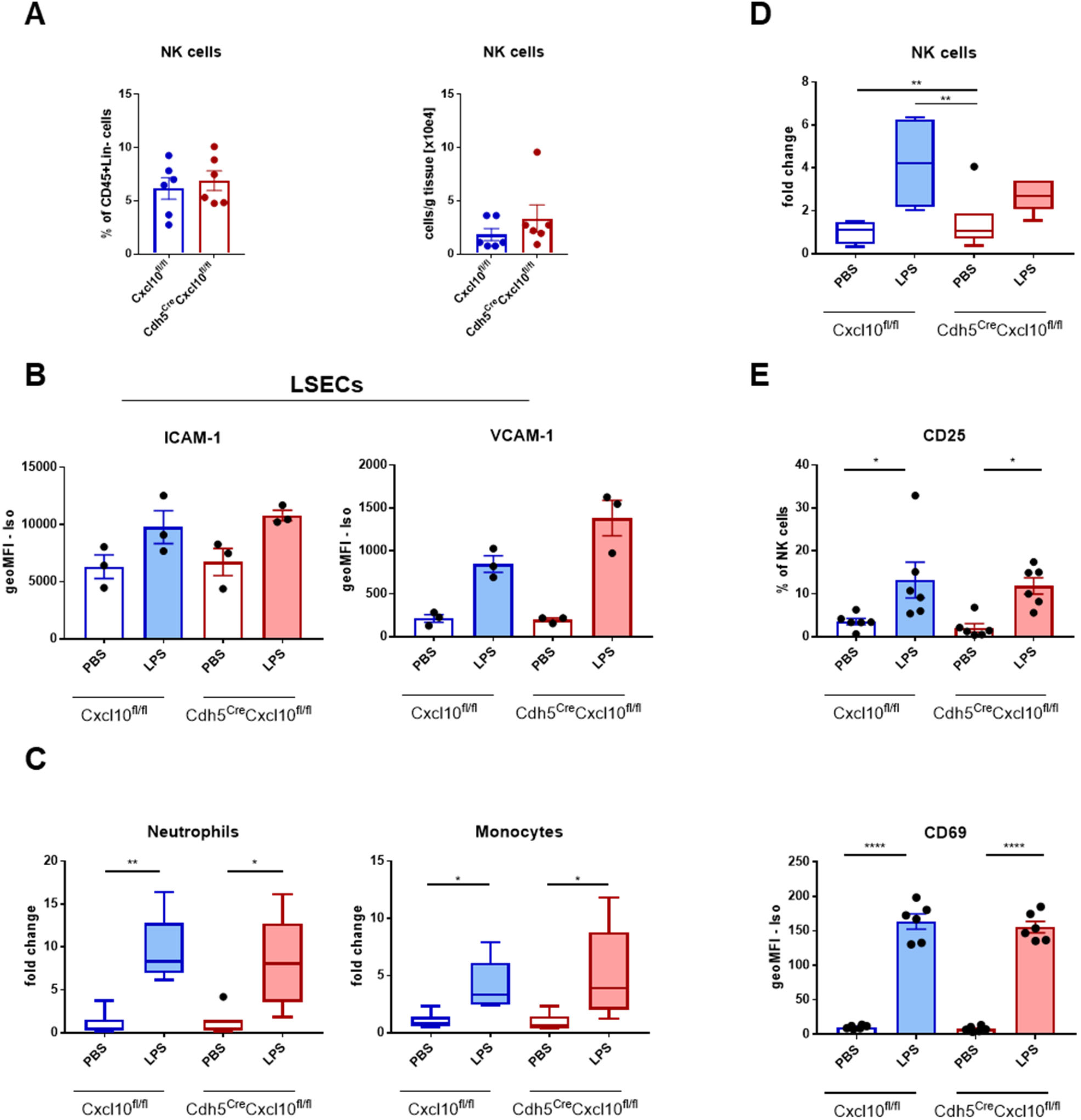
LSEC-derived CXCL10 supports NK cell recruitment to inflamed liver tissue. Cxcl10^fl/fl^ or Cdh5^Cre^Cxcl10^fl/fl^ mice were injected with PBS or LPS, and analyzed 16 h post-injection. (A) Frequencies (left) and cell numbers (right) of NK cells in the livers of Cxcl10^fl/fl^ or Cdh5^Cre^Cxcl10^fl/fl^ mice in steady-state. (B) Expression of ICAM-1 and VCAM-1 on LSECs isolated from PBS- and LPS-injected Cxcl10^fl/fl^ or Cdh5^Cre^Cxcl10^fl/fl^ mice. (C) Fold-change of Neutrophil and Monocyte cell numbers in livers of Cxcl10^fl/fl^ or Cdh5^Cre^Cxcl10^fl/fl^ mice treated with LPS relative to PBS-injected mice. (D) Fold-change of NK cell numbers in the liver tissue. (E) Frequency of CD25-expressing NK cells (top) and CD69 (bottom) expression by NK cells in the livers. (A-B, E) Data are shown as mean ± SEM with each dot representing an individual mouse; n = 6 (A, E) and n = 3 (B). (C-D) In boxplots, box edges represent quartiles, and whiskers represent minimum and maximum data points no more than 1.5 times the interquartile range and line indicates median. Dot denotes outlier. Data are analysed by unpaired ordinary one-way ANOVA. *, p<0.05, **, p<0.01. Iso, isotype; geoMFI, geometric mean fluorescent intensity.

## Discussion

The liver is the primary organ that clears LPS from the bloodstream (Jirillo et al., 2002; Shao et al., 2007; Yao et al., 2016). In steady-state, gut-derived LPS is cleared without inducing tissue inflammation. However, upon infection, increased concentrations of LPS cause activation of both immune and non-parenchymal cells, targeted to remove invading pathogens. Here, we show that in inflammatory settings, LSECs respond to both LPS and immune-derived IFN-γ by modifying their chemokine expression pattern, supporting immune cell recruitment and activation in the liver. TLR4 is expressed by both LSECs and myeloid cells in the liver (Nakamoto and Kanai, 2014; Uhrig et al., 2005). Indeed, we show that LSECs can directly respond to LPS stimulation by increasing the transcription of chemokine genes, such as *Ccl2, Ccl4, Ccl5, Ccl7* and *Ccl2*. Of note, CCL2 was shown to contribute to the recruitment of inflammatory monocytes to the liver (Gschwandtner et al., 2019). The genetic deletion of *Ccl2* led to a reduced accumulation of inflammatory monocytes in a carbon tetrachloride (CCl_4_)-induced liver fibrosis model, and reduced inflammatory damage of the liver (Mitchell et al., 2009). In addition, *Ccr2*-deficient mice displayed reduced liver damage in an acetaminophen-induced acute liver injury model and a non-alcoholic steatohepatitis model (Miura et al., 2012; Mossanen et al., 2016). Therefore, early sensing of high concentrations of LPS by LSECs might provide initial signals for monocyte recruitment, which initiates a downstream inflammatory cascade.

LPS is a major ligand for TLR4 (Akira et al., 2006). Although it was reported that NK cells expressed TLR4 intracellularly (Souza-Fonseca-Guimaraes et al., 2012), studies showed that LPS did not directly act on NK cells to induce IFN-γ production (Varma et al., 2002). The ability of NK cells to increase inflammation by producing IFN-γ was the main mechanism entailing NK cells in LPS-induced mortality (Chan et al., 2014; Emoto et al., 2002; Heremans et al., 1994). Exposure of NK cells to cytokines (Ni et al., 2016) and viral infections (Lau et al., 2018; Luetke-Eversloh et al., 2014) were shown to remodel chromatin and to result in an enhanced IFN-γ production upon re-stimulation. Thus, it is likely that interaction with LSECs led to NK cells acquiring both transcriptomic and epigenetic modifications resulting in an increased IFN-γ production upon stimulation with IL-12 and IL-18.

It was demonstrated that IFN-γ supports chemokine production, including CXCL9, CXCL10, and CXCL11 (Cole et al., 1998; Farber, 1990; Luster et al., 1985). CXCL9, CXCL10 and CXCL11 were shown to contribute to an accumulation of activated lymphocytes, such as NK cells and effector T cells, in inflamed and transformed tissues (Bugide et al., 2021; Dufour et al., 2002; Wendel et al., 2008). We show here that while LPS stimulation directly triggered transcription of the myeloid-recruiting chemokine *Ccl2* by LSECs, the induction of *Cxcl9/10/11* required exposure to IFN-γ. Upon LPS-injection, serum concentration of CXCL10 increased significantly, which correlated with increased accumulation of NK cells in the liver. Conditional deletion of *Cxcl10* in endothelial cells led to a decreased NK cell attraction to the liver, indicating a non-redundant role of LSEC-derived CXCL10 in recruiting NK cells during LPS-induced endotoxemia. The accumulation of NK cells in livers coincided with a decrease of NK cell number in spleens, indicating NK cell recruitment into livers through the portal vein during LPS-induced acute liver injury. IFN-γ was shown to upregulate the expression of adhesion molecules, such as ICAM-1 and VCAM-1 (Ren et al., 2010), which were detected on activated liver endothelium upon challenge with LPS. Therefore, lymphocyte-derived IFN-γ provides a feedback mechanism that supports immune cell recruitment via LSECs, by upregulating both chemokine and adhesion molecule expression on inflamed endothelium.

In summary, we reveal that LSEC-derived chemokines support the accumulation of immune cells, including NK cells and monocytes, in the inflamed liver tissue during an LPS-induced acute liver injury. IFN-γ, but not LPS, which is mainly produced by NK cells and ILC1s in the liver, is required to induce CXCL10 expression by LSECs. In turn, LSECs support IFN-γ production by NK cells, by reshaping their gene expression profile, and by priming enhanced response to myeloid-derived IL-12 and IL-18. Hence, our study highlights LSECs as the non-conventional immune affiliate that orchestrates innate immune cell trafficking and functions during acute liver inflammation.

## Acknowledgments

We thank Petra Bugert, Marian Wincher and Patrick Matei for technical support, the animal facility of the Medical Faculty Mannheim and the German Cancer Research Center for assistance with animal care and experiments, and the ZMF Mannheim for supporting the processing of biological samples.

## Funding

The project was supported by grants from the German Research Foundation [SFB1366 (Project number 319 394046768-SFB 1366; C02 to AC, C01 to MP, B02 to PSRK and Z2 to CM), SPP1937 (CE 140/2-2 to AS and AC), TRR179 (TP07 to AC), SFB-TRR156 (B10N to AC), RTG2099 (Project number: 321 259332240 - RTG2099; P9 to AC and P5 to PSRK), RTG2727 – 445549683 Innate Immune checkpoints in cancer and tissue damage (B1.1 to MP; B1.2 to AC and AS)]; by a network grant of the European Commission (H2020-MSCA-MC322, ITN-765104-MATURE-NK); by the ExU 6.1.11 (to A.C.); by the DKFZ-MOST program (project number 2526) to MP; and by the Angioformatics platform of the European Center for Angioscience (ECAS).

## Author contributions

SP, JXS and AS designed and performed experiments; MJ performed experiments; SP, JXS and AS analyzed the data; CM performed histopathological examination and scoring; SP, JXS, AS and AC wrote the manuscript; CDLT performed gene expression analysis; PSRK provided endothelial cell expertise and discussed the data; AB and MP generated and provided Cdh5^Cre^Cxcl10^fl/fl^ mice and contributed to data discussion; AC and AS designed and supervised the study.

## Declaration of interests

The authors declare no competing interests.

## STAR Methods

### RESOURCE AVAILABILITY

#### Lead contact

Further information and requests for resources and reagents should be directed to and will be fulfilled by the lead contact and/or corresponding authors.

#### Materials Availability

This study did not generate new unique agents.

#### Data and code availability

Gene expression data will be deposited and are publicly available as of the date of publication. Accession numbers will be listed in the key resources table upon publication.

This paper does not report original code.

Any additional information required to analyze the data reported in this paper is available from the lead contact upon request.

### EXPERIMENTAL MODEL AND SUBJECT DETAILS

#### Animals

C57BL6/N wild-type (WT) mice were purchased from Janvier. Mice were housed at the Medical Faculty Mannheim, under specific pathogen-free conditions, and in accordance with all standards of animal care. The Cdh5^Cre^CXCL10^fl/fl^ mice were bred and maintained in the animal facility of the German Cancer Research Center (DKFZ), Heidelberg. All experiments were performed on mice between the age of 10 and 25 weeks. For the LPS injections, male mice were used. For *in vitro* experiments, female mice-derived material was used. All animal experiments were approved by the “Regierungspräsidium Karlsruhe”.

#### LPS-induced acute liver inflammation

Male mice were injected intraperitoneally (i.p.) with LPS (Escherichia coli strain O26:B6, Sigma; 0.5 mg/kg of body weight) re-suspended in PBS. Mice were analyzed 16-18 h upon injection of LPS. Chemokine concentration in serum was assessed using bead-based assay (LEGENDplex™ kit, Biolegend).

### METHODS DETAILS

#### Immune cell isolation and culture

Single-cell suspensions from spleens were prepared by mechanical tissue disruption followed by red blood cell lysis using buffered ammonium chloride potassium phosphate solution. Livers were perfused with PBS, and mechanically and enzymatically digested using Liver Dissociation Kit and gentleMACS™ Octo Dissociator (Miltenyi), according to the manufacturer’s protocol. Immune cells were enriched by density gradient centrifugation over Lympholyte-M (Cedarlane Laboratories) or Percoll gradient (GE Healthcare). NK cells were isolated by flow cytometric sort, and cultured in RPMI-1640 (Gibco) supplemented with 10% FCS, 1% Penicillin/Streptomycin, 1% L-glutamine, 1% MEM non-essential amino acids, 1mM sodium pyruvate, 50mM β-mercaptoethanol (all from Gibco) and 300 U/ml of recombinant human IL-2 (NIH). When indicated, NK cells were stimulated with IL-12 (0.04 ng/ml) and IL-18 (0.8 ng/ml).

#### LSEC isolation and culture

LSECs were purified from dissociated liver tissue by density gradient centrifugation over 26 % Nycodenz (Axis-Shield), followed by magnetic separation with anti-CD146 microbeads (Miltenyi). Purified LSECs (0.8-1×10e6 cells/ml) were cultured on collagen-coated plates (Gibco) in Dulbeccos modified Eagles medium (DMEM, Sigma) supplemented with 10% FCS, 1% Penicillin/Streptomycin, 1% L-glutamine, 1% MEM non-essential amino acids, 1mM sodium pyruvate and 50mM β-mercaptoethanol (all from Gibco). 24 h upon plating, LSECs were stimulated with IFN-γ (100 ng/ml) or LPS (100 ng/ml) overnight. Afterwards, cells were detached with Detachin (Genlantis) and used for staining and flow-cytometry analyses, or lyzed for RNA extraction. For co-culture with NK cells, the LSEC layer was washed with PBS 48 h upon plating, and sorted NK cells (1,25×10e6 cells/ml) were added. Cells were co-cultured in the presence of 300 U/ml of recombinant human IL-2 (NIH) overnight.

#### Flow cytometry and cell sorting

Single-cell suspensions were incubated with FcR-blocking reagent (10% supernatant of αCD16/CD32-producing hybridoma 2.4G2), followed by incubation with fluorochrome-labeled monoclonal antibodies (Key Resources Table). For staining of intracellular antigens, the Intracellular Fixation & Permeabilization Buffer Set (ThermoFisher) was used according to the manufacturer’s instructions. Dead cells were excluded by labeling with 7-aminoactinomycin D (7-AAD, Biolegend), or with ZombieAqua™ (Biolegend,) according to the manufacturer’s instructions. Flow-cytometric analysis or cell sort were conducted with LSRFortessa™ and/or FACSAria™ Fusion flow cytometer (BD Biosciences), respectively. Cells were sorted as following (Supplementary Figure 2): CD45^+^ immune cells, including NKp46-expressing innate lymphocytes (CD3ε^neg^NK1.1^+^NKp46^+^), NK cells (CD3ε^neg^NK1.1^+^, NKp46^+^CD200R^neg^CXCR6^neg^), T cells (Ly6G^neg^CD19^neg^CD3ε^+^TCRβ^+^) Monocytes/Macrophages (Ly6G^neg^CD19^neg^CD11b^+^F4/80^+^), and CD45^neg^ LSECs (CD146^+^CD31^+^Stab2^+^). Data were analyzed using FlowJo™ v10.7.1 Software (BD Life Sciences).

#### Real-time PCR analysis

Sorted cells were lysed and RNA was purified using RNeasy Mini Kit (Qiagen). A total of 400 ng RNA was reverse-transcribed, and cDNA was obtained using the RT^2^ First Strand Kit (Qiagen). For assessment of relative mRNA expression, the mouse Cytokines & Chemokines RT^2^ Profiler array (Qiagen) was used according to the manufacturer’s instructions.

#### Gene Expression Analysis

NK cells isolated from the liver using flow cytometric sort (gated as live CD45^+^CD3^neg^NK1.1^+^NKp46^+^CD200R^neg^TRAIL^neg^ cells) were co-cultured with purified LSECs for 14-16 hours, and then sorted (> 99% of live CD45^+^CD3^neg^NK1.1^+^NKp46^+^ cells) for gene expression analysis. RNA was isolated using RNeasy Mini Kit (Qiagen), and genomic DNA was removed by TURBO™-DNase (Thermo Fisher Scientific) according to the manufacturer’s protocol. Gene expression profiling was performed using arrays of Clariom™ D Arrays (Thermo Fisher Scientific). Biotinylated antisense cDNA was prepared according to the standard labeling protocol with the GeneChip® WT Plus Reagent Kit and the GeneChip® Hybridization, Wash and Stain Kit (both from Thermo Fisher Scientific). Thereafter, chip hybridization was performed on a GeneChip Hybridization oven 640, then dyed in the GeneChip Fluidics Station 450, and afterwards scanned with a GeneChip Scanner 3000. All of the equipment used was from the Affymetrix-Company (Affymetrix, High Wycombe, UK). Arrays raw fluorescence intensity values were normalized by applying quantile normalization and RMA background correction. A custom CDF version 22 with ENTREZ based gene definitions was used to annotate the arrays (Dai et al., 2005). Differentially expressed genes were obtained by means of OneWay-ANOVA using a commercial software package SAS JMP15 Genomics, version 10, from SAS (SAS Institute, Cary, NC, USA). Parameters applied were a false positive rate of α = 0.05 with FDR correction as the level of significance. Data are visualized using *EnhancedVolcano* package in R (https://github.com/kevinblighe/EnhancedVolcano). Upregulated gene transcripts (p-value < 0.0001 and FC > 1.2) were submitted to evaluate the enrichment of GO terms (GO, http://geneontology.org/) using the *clusterProfiler* package (Wu et al., 2021).

### QUANTIFICATION AND STATISTICAL ANALYSIS

For statistical analyses and data visualization, Prism 7.03 (Graphpad), R (4.1x and 4.2x), RStudio (1.4.1717), and packages *ggplot2* (Wickham, 2016) and *pheatmap* (https://www.rdocumentation.org/packages/pheatmap/versions/0.2/topics/pheatmap) were used. Data are tested for normal distribution using Shapiro-Wilk test followed by evaluation using an appropriate test, as indicated in Figure Legends. Correction for multiple comparison testing was done when necessary. Experimental results are shown as mean ± SEM; n represents numbers of animals or cells in the experiments, as specified in Figure Legends. Experimental groups were considered to be significantly different when *, p<0.05, **, p<0.005, ***, p<0.001 and ****, p<0.0001.

## KEY RESOURCES TABLE

| REAGENT or RESOURCE | SOURCE | IDENTIFIER |
| --- | --- | --- |
| **Antibodies** |  |  |
| Anti-mouse Arm Hamster IgG FITC (HTK888) | Biolegend | Cat#400906 |
| Anti-mouse CD3 APC-Cy7 (14A2) | Biolegend | Cat#100222<br>RRID:AB_2242784 |
| Anti-mouse CD3 BV711 (145-2C11) | Biolegend | Cat#100349<br>RRID:AB_2565841 |
| Anti-mouse CD3 PE-Cy7 (145-2C11) | Biolegend | Cat#100320<br>RRID:AB_312685 |
| Anti-mouse CD11b APC (M1/70) | Biolegend | Cat#101212 |
| Anti-mouse/human CD11b BV650 (M1/70) | Biolegend | Cat#101259;<br>RRID:AB_2566568 |
| Anti-mouse CD11c PE-Cy7 (N418) | Biolegend | Cat#117318<br>RRID:AB_312795 |
| Anti-mouse CD19 APC-Cy7 (6D5) | Biolegend | Cat#115530<br>RRID:AB_830707 |
| Anti-mouse CD25 PE (PC61) | Biolegend | Cat#102008<br>RRID:AB_312857 |
| Anti-mouse CD31 BV421 (390) | Biolegend | Cat#102424<br>RRID:AB_2650892 |
| Anti-mouse CD45 BV785 (30F11) | Biolegend | Cat#103149<br>RRID:AB_2564590 |
| Anti-mouse CD45 PerCP-Cy5.5 (30F11) | Biolegend | Cat#103132<br>RRID:AB_893340 |
| Anti-mouse CD69 FITC (H1.2F3) | Biolegend | Cat#104506<br>RRID:AB_313109 |
| Anti-mouse CD146 PE-Cy7 (ME-9F1) | Biolegend | Cat#134714<br>RRID:AB_2563109 |
| Anti-mouse CXCR3 BV421 (CXCR3-173) | Biolegend | Cat#126529<br>RRID:AB_2563100 |
| Anti-mouse CXCR3 PE-Cy7 (CXCR3-173) | Biolegend | Cat#126516<br>RRID:AB_2245493 |
| Anti-mouse Eomes PE-Cy7 (Dan-11mag) | Thermo Fisher Scientific | Cat#25-4875-82<br>RRID:AB_2573454 |
| Anti-mouse Eomes TexasRed (X4-83) | BD Horizon | Cat#567167<br>RRID:AB_2916484 |
| Anti-mouse F4/80 APC-Cy7 (BM8) | Biolegend | Cat#123118<br>RRID:AB_893477 |
| Anti-mouse F4/80 PE/Dazzle 594 (BM8) | Biolegend | Cat#123146<br>RRID:AB_2564133 |
| Anti-mouse ICAM-1 Alexa Flour 647 (YNI/1.7.4) | Biolegend | Cat#116114<br>RRID:AB_493495 |
| Anti-mouse IFNg Alexa Flour 647 (XMG1.2) | Biolegend | Cat#505814<br>RRID:AB_493314 |
| Anti-mouse Ki67 PE (SolA15) | eBiosciences | 12-5698-82 |
| Anti-mouse Ly6C FITC (HK1.4) | Biolegend | Cat#128005<br>RRID:AB_1186134 |
| Anti-mouse Ly6G APC-Cy7 (1A8) | Biolegend | Cat#127624<br>RRID:AB_10640819 |
| Anti-mouse Ly6G Pacific Blue (1A8) | Biolegend | Cat#127612<br>RRID:AB_2251161 |
| Anti-mouse NK1.1 BV785 (PK136) | Biolegend | Cat#108749<br>RRID:AB_2564304 |
| Anti-mouse NK1.1 PE (PK136) | Biolegend | Cat#108708<br>RRID:AB_313395 |
| Anti-mouse NKp46 APC (19A1.4) | Biolegend | Cat#137608<br>RRID:AB_10612758 |
| Anti-mouse NKp46 BV421 (27A1.4) | Biolegend | Cat#137612<br>RRID:AB_2563104 |
| Anti-mouse Rat IgG1 PE (G0114F7) | Biolegend | Cat#401906 |
| Anti-mouse Rat IgG2a APC (54447) | R&D Systems | Cat#IC006A<br>RRID:AB_357254 |
| Anti-mouse Rat IgG2a BV421 (RTK2758) | Biolegend | Cat#400549<br>RRID:AB_2885080 |
| Anti-mouse Siglec F APC-Cy7 (E50-2440) | Biolegend | Cat#565527 |
| Anti-mouse Stabilin-2 FITC (34-2) | MBL International | Cat#D317-A48<br>RRID:AB_2811155 |
| Anti-mouse TCRb BV785 (H57-597) | Biolegend | Cat#109249<br>RRID:AB_2810347 |
| Anti-mouse Ter119 APC-Cy7 (Ter-119) | Biolegend | Cat#116223<br>RRID:AB_2137788 |
| Anti-mouse VCAM-1 APC (429(MVCAM.A)) | Biolegend | Cat#105712<br>RRID:AB_493429 |
| **Chemicals, peptides, and recombinant proteins** |  |  |
| Lipopolysaccharide from E. coli (O26:B6) | Sigma | Cat#L2654;<br>CAS: 93572-42-0 |
| Recombinant human IL-2 | Hoffmann-La Roche | Cat#1104-0890 |
| Recombinant mouse IL-12 | Peprtech | Cat#210-12 |
| Recombinant mouse IL-18 | MBL | Cat#B002-5 |
| Recombinant mouse IFN-γ | Peprtech | Cat#315-05 |
| BD Golgi Plug | BD Biosciences | Cat#555029 |
| 7-AAD | BD Biosciences | Cat#420404 |
| Zombie Aqua Fixable Viability Dye | Biolegend | Cat#423102 |
| Nycodenz | Axis-Shield | Cat#1002424 |
| Percoll | GE Healthcare | Cat#17089101 |
| Lympholyte M | Cedarlane | Cat#CL5035 |
| **Critical commercial assays** |  |  |
| Liver Dissociation Kit Mouse | Mitlenyi | Cat#130-105-807 |
| FoxP3 Transcription factor staining buffer set | eBioscience | Cat#00-5523-00 |
| LEGENDplex Mouse | Biolegend | Cat#740451,<br>Cat#741068 |
| RT2 Profiler PCR Array Mouse Cytokines & Chemokines (PAMM-150ZE-4) | QIAGEN | Cat#330231 |
| Clariom™ D Array, mouse | Affymetrix | Cat#902512 |
| **Experimental models: Organisms/strains** |  |  |
| Mouse: C57BL/6N (C57BL/6NRj) | Janvier Labs | RRID:MGI:6236253 |
| Mouse: B6-Cxcl10tm1.1Dple Tg(Cdh5-cre)1Spe/Platt (Cdh5<sup>Cre</sup>Cxcl10<sup>fl/fl</sup>) | DKFZ, Heidelberg | N/A |
| **Software and algorithms** |  |  |
| FlowJo version >10.7.1 | FlowJo LLC | https://www.flowjo.com/ |
| GraphPad Prism 7 | GraphPad | https://www.graphpad.com/ |
| LEGENDplex Data Analysis Software | Biolegend | https://www.biolegend.com/en-us/legendplex |
| R version >4.1 | R core team | https://www.r-project.org/ |
| **Other** |  |  |
| Collagen I coated well plates | Gibco | Cat#11142802 |
| Gentle MACS C tubes | Miltenyi | Cat#130-093-237 |
| MACS LS Columns | Miltenyi | Cat#130-042-401 |
| CD146 (LSEC) microbeads, mouse | Miltenyi | Cat#130-092-007 |

## Notes

### Competing Interest Statement

The authors have declared no competing interest.

## References

Adams, N.M., Lau, C.M., Fan, X., Rapp, M., Geary, C.D., Weizman, O.E., Diaz-Salazar, C., and Sun, J.C. (2018). Transcription Factor IRF8 Orchestrates the Adaptive Natural Killer Cell Response. Immunity 48, 1172–1182 e1176.

Akira, S., Uematsu, S., and Takeuchi, O. (2006). Pathogen recognition and innate immunity. Cell 124, 783–801.

Amersfoort, J., Eelen, G., and Carmeliet, P. (2022). Immunomodulation by endothelial cells - partnering up with the immune system? Nat Rev Immunol.

Beaulieu, A.M., Zawislak, C.L., Nakayama, T., and Sun, J.C. (2014). The transcription factor Zbtb32 controls the proliferative burst of virus-specific natural killer cells responding to infection. Nature Immunology 15, 546–553.

Bugide, S., Gupta, R., Green, M.R., and Wajapeyee, N. (2021). EZH2 inhibits NK cell-mediated antitumor immunity by suppressing CXCL10 expression in an HDAC10-dependent manner. Proc Natl Acad Sci U S A 118.

Cabral, F., Al-Rahem, M., Skaggs, J., Thomas, T.A., Kumar, N., Wu, Q., Fadda, P., Yu, L., Robinson, J.M., Kim, J., et al. (2021). Stabilin receptors clear LPS and control systemic inflammation. iScience 24, 103337.

Chan, C.J., Martinet, L., Gilfillan, S., Souza-Fonseca-Guimaraes, F., Chow, M.T., Town, L., Ritchie, D.S., Colonna, M., Andrews, D.M., and Smyth, M.J. (2014). The receptors CD96 and CD226 oppose each other in the regulation of natural killer cell functions. Nat Immunol 15, 431–438.

Cole, K.E., Strick, C.A., Paradis, T.J., Ogborne, K.T., Loetscher, M., Gladue, R.P., Lin, W., Boyd, J.G., Moser, B., Wood, D.E., et al. (1998). Interferon-inducible T cell alpha chemoattractant (I-TAC): a novel non-ELR CXC chemokine with potent activity on activated T cells through selective high affinity binding to CXCR3. J Exp Med 187, 2009–2021.

Dai, M., Wang, P., Boyd, A.D., Kostov, G., Athey, B., Jones, E.G., Bunney, W.E., Myers, R.M., Speed, T.P., and Akil, H. (2005). Evolving gene/transcript definitions significantly alter the interpretation of GeneChip data. Nucleic acids research 33, e175–e175.

Ducimetiere, L., Lucchiari, G., Litscher, G., Nater, M., Heeb, L., Nunez, N.G., Wyss, L., Burri, D., Vermeer, M., Gschwend, J., et al. (2021). Conventional NK cells and tissue-resident ILC1s join forces to control liver metastasis. Proc Natl Acad Sci U S A 118.

Dufour, J.H., Dziejman, M., Liu, M.T., Leung, J.H., Lane, T.E., and Luster, A.D. (2002). IFN-gamma-inducible protein 10 (IP-10; CXCL10)-deficient mice reveal a role for IP-10 in effector T cell generation and trafficking. J Immunol 168, 3195–3204.

Emoto, M., Miyamoto, M., Yoshizawa, I., Emoto, Y., Schaible, U.E., Kita, E., and Kaufmann, S.H. (2002). Critical role of NK cells rather than V alpha 14(+)NKT cells in lipopolysaccharide-induced lethal shock in mice. J Immunol 169, 1426–1432.

Farber, J.M. (1990). A macrophage mRNA selectively induced by gamma-interferon encodes a member of the platelet factor 4 family of cytokines. Proc Natl Acad Sci U S A 87, 5238–5242.

Fehniger, T.A., Shah, M.H., Turner, M.J., VanDeusen, J.B., Whitman, S.P., Cooper, M.A., Suzuki, K., Wechser, M., Goodsaid, F., and Caligiuri, M.A. (1999). Differential cytokine and chemokine gene expression by human NK cells following activation with IL-18 or IL-15 in combination with IL-12: Implications for the innate immune response. Journal of Immunology 162, 4511–4520.

Ganesan, L.P., Kim, J., Wu, Y., Mohanty, S., Phillips, G.S., Birmingham, D.J., Robinson, J.M., and Anderson, C.L. (2012). FcgammaRIIb on liver sinusoidal endothelium clears small immune complexes. J Immunol 189, 4981–4988.

Ganesan, L.P., Mohanty, S., Kim, J., Clark, K.R., Robinson, J.M., and Anderson, C.L. (2011). Rapid and efficient clearance of blood-borne virus by liver sinusoidal endothelium. Plos Pathog 7, e1002281.

Geraud, C., Koch, P.S., Zierow, J., Klapproth, K., Busch, K., Olsavszky, V., Leibing, T., Demory, A., Ulbrich, F., Diett, M., et al. (2017). GATA4-dependent organ-specific endothelial differentiation controls liver development and embryonic hematopoiesis. J Clin Invest 127, 1099–1114.

Geraud, C., Mogler, C., Runge, A., Evdokimov, K., Lu, S., Schledzewski, K., Arnold, B., Hammerling, G., Koch, P.S., Breuhahn, K., et al. (2013). Endothelial transdifferentiation in hepatocellular carcinoma: loss of Stabilin-2 expression in peri-tumourous liver correlates with increased survival. Liver Int 33, 1428–1440.

Gschwandtner, M., Derler, R., and Midwood, K.S. (2019). More Than Just Attractive: How CCL2 Influences Myeloid Cell Behavior Beyond Chemotaxis. Frontiers in Immunology 10.

Harjunpaa, H., Asens, M.L., Guenther, C., and Fagerholm, S.C. (2019). Cell Adhesion Molecules and Their Roles and Regulation in the Immune and Tumor Microenvironment. Frontiers in Immunology 10.

Heremans, H., Dillen, C., van Damme, J., and Billiau, A. (1994). Essential role for natural killer cells in the lethal lipopolysaccharide-induced Shwartzman-like reaction in mice. Eur J Immunol 24, 1155–1160.

Jirillo, E., Caccavo, D., Magrone, T., Piccigallo, E., Amati, L., Lembo, A., Kalis, C., and Gumenscheimer, M. (2002). The role of the liver in the response to LPS: experimental and clinical findings. J Endotoxin Res 8, 319–327.

Knolle, P.A., and Wohlleber, D. (2016). Immunological functions of liver sinusoidal endothelial cells. Cell Mol Immunol 13, 347–353.

Lau, C.M., Adams, N.M., Geary, C.D., Weizman, O.E., Rapp, M., Pritykin, Y., Leslie, C.S., and Sun, J.C. (2018). Epigenetic control of innate and adaptive immune memory. Nat Immunol 19, 963–972.

Loetscher, M., Gerber, B., Loetscher, P., Jones, S.A., Piali, L., Clark-Lewis, I., Baggiolini, M., and Moser, B. (1996). Chemokine receptor specific for IP10 and mig: structure, function, and expression in activated T-lymphocytes. J Exp Med 184, 963–969.

Loftus, R.M., Assmann, N., Kedia-Mehta, N., O’Brien, K.L., Garcia, A., Gillespie, C., Hukelmann, J.L., Oefner, P.J., Lamond, A.I., Gardiner, C.M., et al. (2018). Amino acid-dependent cMyc expression is essential for NK cell metabolic and functional responses in mice. Nature Communications 9.

Long, E.O., Kim, H.S., Liu, D.F., Peterson, M.E., and Rajagopalan, S. (2013). Controlling Natural Killer Cell Responses: Integration of Signals for Activation and Inhibition. Annual Review of Immunology, Vol 31 31, 227–258.

Luetke-Eversloh, M., Hammer, Q., Durek, P., Nordstrom, K., Gasparoni, G., Pink, M., Hamann, A., Walter, J., Chang, H.D., Dong, J., et al. (2014). Human Cytomegalovirus Drives Epigenetic Imprinting of the IFNG Locus in NKG2C(hi) Natural Killer Cells. Plos Pathog 10.

Luster, A.D., and Ravetch, J.V. (1987). Biochemical characterization of a gamma interferon-inducible cytokine (IP-10). J Exp Med 166, 1084–1097.

Luster, A.D., Unkeless, J.C., and Ravetch, J.V. (1985). Gamma-interferon transcriptionally regulates an early-response gene containing homology to platelet proteins. Nature 315, 672–676.

Lusty, E., Poznanski, S.M., Kwofie, K., Mandur, T.S., Lee, D.A., Richards, C.D., and Ashkar, A.A. (2017). IL-18/IL-15/IL-12 synergy induces elevated and prolonged IFN-gamma production by ex vivo expanded NK cells which is not due to enhanced STAT4 activation. Mol Immunol 88, 138–147.

Mitchell, C., Couton, D., Couty, J.P., Anson, M., Crain, A.M., Bizet, V., Renia, L., Pol, S., Mallet, V., and Gilgenkrantz, H. (2009). Dual Role of CCR2 in the Constitution and the Resolution of Liver Fibrosis in Mice. Am J Pathol 174, 1766–1775.

Miura, K., Yang, L., van Rooijen, N., Ohnishi, H., and Seki, E. (2012). Hepatic recruitment of macrophages promotes nonalcoholic steatohepatitis through CCR2. Am J Physiol Gastrointest Liver Physiol 302, G1310–1321.

Mossanen, J.C., Krenkel, O., Ergen, C., Govaere, O., Liepelt, A., Puengel, T., Heymann, F., Kalthoff, S., Lefebvre, E., Eulberg, D., et al. (2016). Chemokine (C-C motif) receptor 2-positive monocytes aggravate the early phase of acetaminophen-induced acute liver injury. Hepatology 64, 1667–1682.

Mujal, A.M., Delconte, R.B., and Sun, J.C. (2021). Natural Killer Cells: From Innate to Adaptive Features. Annu Rev Immunol 39, 417–447.

Nakamoto, N., and Kanai, T. (2014). Role of toll-like receptors in immune activation and tolerance in the liver. Front Immunol 5, 221.

Ni, J., Holsken, O., Miller, M., Hammer, Q., Luetke-Eversloh, M., Romagnani, C., and Cerwenka, A. (2016). Adoptively transferred natural killer cells maintain long-term antitumor activity by epigenetic imprinting and CD4(+) T cell help. Oncoimmunology 5, e1219009.

Ren, G.W., Zhao, X., Zhang, L.Y., Zhang, J.M., L’Huillier, A., Ling, W.F., Roberts, A.I., Le, A.D., Shi, S.T., Shao, C.S., et al. (2010). Inflammatory Cytokine-Induced Intercellular Adhesion Molecule-1 and Vascular Cell Adhesion Molecule-1 in Mesenchymal Stem Cells Are Critical for Immunosuppression. Journal of Immunology 184, 2321–2328.

Shao, B., Lu, M., Katz, S.C., Varley, A.W., Hardwick, J., Rogers, T.E., Ojogun, N., Rockey, D.C., Dematteo, R.P., and Munford, R.S. (2007). A host lipase detoxifies bacterial lipopolysaccharides in the liver and spleen. J Biol Chem 282, 13726–13735.

Souza-Fonseca-Guimaraes, F., Parlato, M., Fitting, C., Cavaillon, J.M., and Adib-Conquy, M. (2012). NK cell tolerance to TLR agonists mediated by regulatory T cells after polymicrobial sepsis. J Immunol 188, 5850–5858.

Sun, X., Wu, J., Liu, L., Chen, Y., Tang, Y., Liu, S., Chen, H., Jiang, Y., Liu, Y., Yuan, H., et al. (2022). Transcriptional switch of hepatocytes initiates macrophage recruitment and T-cell suppression in endotoxemia. J Hepatol.

Uhrig, A., Banafsche, R., Kremer, M., Hegenbarth, S., Hamann, A., Neurath, M., Gerken, G., Limmer, A., and Knolle, P.A. (2005). Development and functional consequences of LPS tolerance in sinusoidal endothelial cells of the liver. J Leukoc Biol 77, 626–633.

Varma, T.K., Lin, C.Y., Toliver-Kinsky, T.E., and Sherwood, E.R. (2002). Endotoxin-induced gamma interferon production: contributing cell types and key regulatory factors. Clin Diagn Lab Immunol 9, 530–543.

Vinay, D.S., Choi, B.K., Bae, J.S., Kim, W.Y., Gebhardt, B.M., and Kwon, B.S. (2004). CD137-deficient mice have reduced NK/NKT cell numbers and function, are resistant to lipopolysaccharide-induced shock syndromes, and have lower IL-4 responses. J Immunol 173, 4218–4229.

Weizman, O.E., Adams, N.M., Schuster, I.S., Krishna, C., Pritykin, Y., Lau, C., Degli-Esposti, M.A., Leslie, C.S., Sun, J.C., and O’Sullivan, T.E. (2017). ILC1 Confer Early Host Protection at Initial Sites of Viral Infection. Cell 171, 795–+.

Wendel, M., Galani, I.E., Suri-Payer, E., and Cerwenka, A. (2008). Natural Killer Cell Accumulation in Tumors Is Dependent on IFN-gamma and CXCR3 Ligands. Cancer Research 68, 8437–8445.

Wickham, H. (2016). ggplot2: Elegant Graphics for Data Analysis. Springer-Verlag New York.

Wu, T., Hu, E., Xu, S., Chen, M., Guo, P., Dai, Z., Feng, T., Zhou, L., Tang, W., Zhan, L., et al. (2021). clusterProfiler 4.0: A universal enrichment tool for interpreting omics data. Innovation (Camb) 2, 100141.

Yao, Z., Mates, J.M., Cheplowitz, A.M., Hammer, L.P., Maiseyeu, A., Phillips, G.S., Wewers, M.D., Rajaram, M.V., Robinson, J.M., Anderson, C.L., et al. (2016). Blood-Borne Lipopolysaccharide Is Rapidly Eliminated by Liver Sinusoidal Endothelial Cells via High-Density Lipoprotein. J Immunol 197, 2390–2399.

