## Supplemental Information for "Liver sinusoidal endothelial cells orchestrate NK cell recruitment and activation in acute inflammatory liver injury"

[1]Department of Immunobiochemistry, Mannheim Institute for Innate Immunoscience (MI3), Medical Faculty Mannheim, Heidelberg University, Mannheim, Germany
[2]European Center for Angioscience (ECAS), Medical Faculty Mannheim, Heidelberg University, Mannheim, Germany
[3]NGS Core Facility, Medical Faculty Mannheim, Heidelberg University, Mannheim, Germany
[4]Department of Dermatology, Venereology and Allergology, University Medical Center and Medical Faculty Mannheim, Heidelberg University, and Center of Excellence in Dermatology, Mannheim, Germany
[5]CCU Neuroimmunology and Brain Tumor Immunology, German Cancer Research Center (DKFZ), German Cancer Consortium (DKTK), Clinical Cooperation Unit (CCU), Heidelberg, Germany
[6]Institute of Pathology, Technical University Munich, Munich, Germany
[7]Department of Neurology, University Hospital Mannheim (UMM), MCTN, Medical Faculty Mannheim, Heidelberg University, Mannheim, Germany
[8]DKFZ Hector Cancer Institute at the University Medical Center Mannheim, Mannheim, Germany
[#]Equally contributing first authors
[*]Equally contributing senior authors

### Supplemental Figure 1

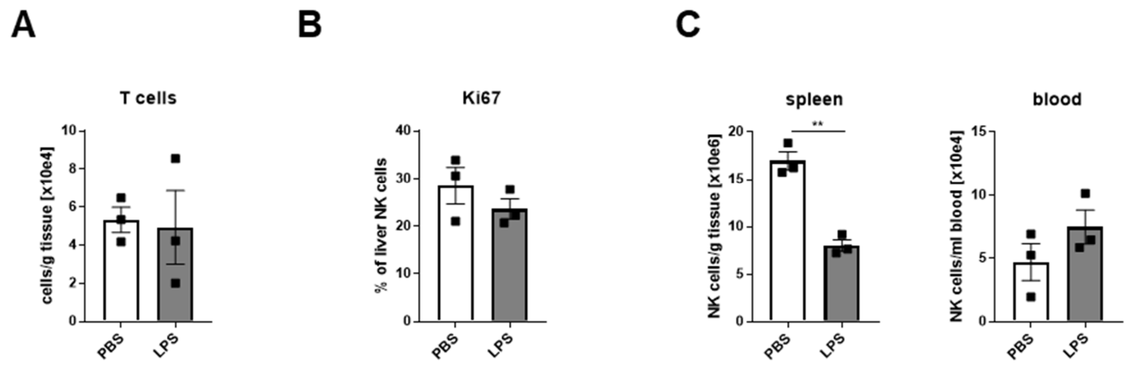


**Supplemental Figure 1. T cell frequency in livers, Ki67 expression by hepatic NK cells, and NK cell numbers in spleen and blood of LPS-treated mice**

(A) T cell numbers (CD45$^+$Lineage(Ly6G, SiglecF, F4/80, CD19)$^{neg}$CD3ε$^+$TCRb$^+$) in the livers of LPS-injected mice. (B) Frequency of Ki67-expressing NK cells in the livers of PBS- or LPS-injected mice. (C) NK cell numbers in the spleen and the blood of PBS- or LPS-challenged mice. (A-C) Data are shown as mean ± SEM, analyzed by unpaired Student's t-Test. Related to Figure 1.

**Supplemental Figure 2**

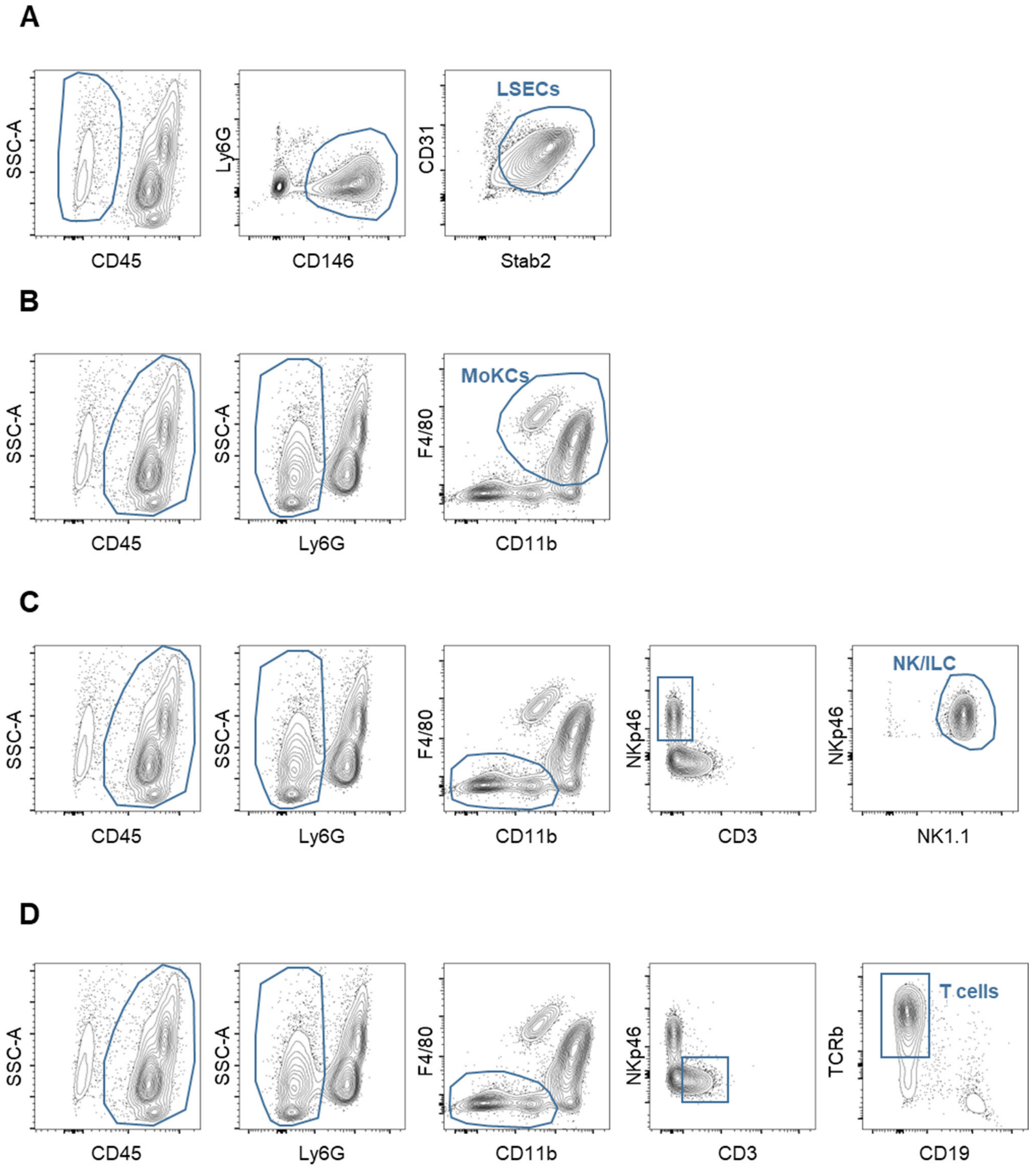

**Supplemental Figure 2. Sorting strategy for LSECs and immune cell subsets from inflamed liver tissue**
(A) Mice were injected with LPS intraperitoneally. Single-cell suspensions were prepared from pooled liver tissue of 4 mice, and LSECs and indicated immune cell subsets were purified by flow-cytometric sort. All subsets were pre-gated as single and alive cells. (A) LSECs were sorted as $CD45^{neg}Ly6G^{neg}CD146^{+}CD31^{+}Stab2^{+}$. (B) Among CD45-expressing cells, Monocytes and Kupffer cells (MoKCs) were sorted as $Ly6G^{neg}F4/80^{+}CD11b^{+}$. (C) From Neutrophil ($Ly6G^{+}$)- and MoKC ($F4/80^{+}$)-excluded fraction, NK cells and ILC1s were purified as $CD3\varepsilon^{neg}NK1.1^{+}NKp46^{+}$. (D) T cells were purified after NK cell exclusion as $CD3\varepsilon^{+}TCRb^{+}CD19^{neg}$. Related to Figure 3 and Figure 5.

**Supplemental Figure 3**

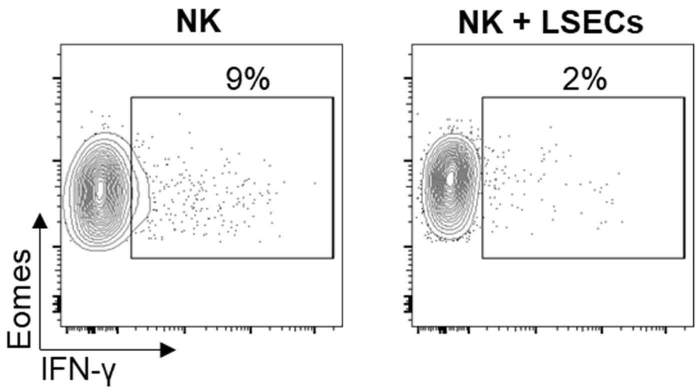


**Supplemental Figure 3. LSECs do not induce IFN-γ production by NK cells**

Representative contour plots of IFN-γ expression by NK cells cultured alone or with LSECs for 16 h.Related to Figure 5.
